# Electrophysiological responses to appetitive and consummatory behavior in the rostral nucleus tractus solitarius in awake, unrestrained rats

**DOI:** 10.1101/2024.04.30.591929

**Authors:** Stephen A. Pilato, Flynn P. O’Connell, Jonathan D. Victor, Patricia M. Di Lorenzo

**Affiliations:** Dept. of Psychology, Box 6000, Binghamton University, Binghamton, NY 13902-6000; Elizabeth R. Miller Brain Observatory, The Rockefeller University, NY, NY; Feil Family Brain and Mind Research Institute, Weill Cornell Medical College, NY, NY 10065

## Abstract

As the intermediate nucleus in the brainstem receiving information from the tongue and transmitting information upstream, the rostral portion of the nucleus tractus solitarius (rNTS) is most often described as a “taste relay”. Although recent evidence implicates the NTS in a broad neural circuit involved in regulating ingestion, there is little information about how cells in this structure respond when an animal is eating solid food. Here, single cells in the rNTS were recorded in awake, unrestrained rats as they explored and ate solid foods (Eating paradigm) chosen to correspond to the basic taste qualities: milk chocolate for sweet, salted peanuts for salty, Granny Smith apples for sour and broccoli for bitter. A subset of cells was also recorded as the animal licked exemplars of the five basic taste qualities: sucrose, NaCl, citric acid, quinine and MSG (Lick paradigm). Results showed that most cells were excited by exploration of a food-filled well, sometimes responding prior to contact with the food. In contrast, cells that were excited by food well exploration became significantly less active while the animal was eating the food. Most cells were broadly tuned across foods, and those cells that were recorded in both the Lick and Eating paradigms showed little correspondence in their tuning across paradigms. The preponderance of robust responses to the appetitive versus the consummatory phase of ingestion suggests that multimodal convergence onto cells in the rNTS may be used in decision making about ingestion.

**Significance Statement:** The rostral part of the NTS has traditionally, but perhaps narrowly, been thought of as a “taste relay”. While it is true that this structure receives and transmits information about tastants in the mouth to higher order structures in the central gustatory pathway, data presented here show that its function is more diverse. Specifically, in addition to the responses to liquid tastants in the mouth, its responses to approach and exploration of solid food define a significant role for the rNTS in the appetitive phase of eating. Moreover, responses to food consumption, albeit weaker than those during appetitive behavior, buttress the idea that the rNTS is part of the larger hindbrain circuit guiding energy regulation.

## Introduction

It has been argued that the neural circuits that guide the regulation of food intake begin in the brainstem (Cheng et al., 2022; Grill and Hayes, 2012), specifically in the intermediate and caudal nucleus tractus solitarius (NTS). However, the contribution of the rostral, gustatory NTS (rNTS) to this function has not been explored, except as a footnote for providing sensory and hedonic evaluation of ingesta. Along with the more caudal NTS, the extensive and parallel connections of the rNTS to both rostral and caudal (King, 2007) structures involved with ingestion suggest that the rNTS may play a more significant role in feeding behavior than traditionally conceptualized.

The rNTS is the first node in the central gustatory neural pathway (Vincis and Fontanini, 2019). While it is often referred to as a simple “relay” in the taste system, it is the site of considerable integration of sensory inputs along with taste, including olfactory (Escanilla and Di Lorenzo, 2015), somatosensory (Ogawa and Imoto, 1984) and thermal (Wilson and Lemon, 2013) senses. In addition, the firing patterns of a significant proportion of rNTS cells in the alert, freely licking animal have been shown to reflect motor output (Roussin et al., 2012; Denman et al., 2019). Thus, it can be argued that the rNTS is multimodal as well as sensorimotor in character (Di Lorenzo, 2021), advancing the idea that it is an active participant in the feeding process.

In spite of the evidence that the rNTS is multimodal in its sensitivity profile, the preponderance of research on the rNTS has been narrowly focused on the responses to taste in its most reductionist form. That is, decades of studies have reported neural responses in rNTS to exemplars of five prototypical taste stimuli representing the five so-called “basic” taste qualities: sweet, salty, sour, bitter and umami (savory). Theories about how taste is represented in the rNTS have been most commonly based on neurophysiological recordings from cells in anesthetized preparations. Given the more recent findings suggesting that cells in the rNTS reflect a broad multimodal repertoire, it is reasonable to suggest that the function of the rNTS is to represent information about food and feeding, rather than purely taste as it has been traditionally conceived.

If indeed the gustatory system is purposed for identifying the sensory aspects of food, broadly defined, then investigations about what the gustatory system is doing while an animal is eating food are necessary – but largely lacking. One exception is a study by Yamamoto et al. (1988). In that study, responses of gustatory cortical neurons were recorded while subjects were licking tastants, grooming, and eating food pellets. A subset of taste-responsive neurons that became active when the rat ate food was identified. Yamamoto et al. (1988) argued that these food-related responses were a reflection of mechanosensation. Since the food pellets were not flavored, it was not possible to evaluate what role, if any, the taste quality of the food affected eating behavior. Interestingly, Yamamoto et al. (1988) also found cells that were responsive to olfactory stimuli as well as others that were anticipatory with respect to licking tastants. There were also taste-responsive neurons that decreased their firing rates in response to palatable stimuli and to food pellets, a result that Yamamoto et al. (1988) attributed to familiarity with the food pellet.

Here, we recorded rNTS responses in awake, unrestrained rats as they approached and consumed various solid foods and as they licked traditional tastants. Tastants included exemplars of the five basic taste qualities: sweet, salty, sour, bitter and umami. Solid foods were chosen such that the predominant taste was one of these basic taste qualities. Results showed that cells and the rNTS were particularly responsive to the approach and sampling of solid food but less responsive when these foods were being eaten. In all, these data suggest a reconceptualization of the functionality of the rNTS with respect to feeding behavior.

## Materials and Methods

### Subjects

Subjects were male (*n* = 7) and female (*n* = 1) Sprague Dawley rats (Taconic labs) weighing ranged 250 – 750g and maintained 12:12-h light-dark cycle (lights off at 0900 hours). Rats were tested during the first 6 hours of the dark period. Animals were pair-housed in plastic cages with environmental enrichment until electrode implantation after which they were single-housed. Standard chow was available *ad libitum*. During data collection, water was available for one hour daily in addition to fluid consumption during the experimental paradigm. Animal care and procedures were approved by the Binghamton University Institutional Animal Care and Use Committee.

### Apparatus

A clear Plexiglas experimental chamber (Med Associates, Fairfax, VT) housed in a melamine box was used for data collection. A window, also made of clear Plexiglas, was located on the front door of the melamine box which allowed for observation and video recording of the rat’s behavior. A stainless steel sipper tube was positioned just behind an opening on one wall of the experimental chamber. Licks were recorded when an animal broke an infrared photobeam across the opening. The sipper tube housed a collection of smaller stainless steel tubes that were each fed by a separate pressurized reservoir of a taste stimulus. A computer-activated solenoid was interposed between the tastant reservoir and the sipper tube. Each time the rat licked, except for “dry licks,” 12 µl ± 2 µl of fluid was delivered. This arrangement allowed the experimenter to control what stimulus the rat was given on a lick-by-lick basis. Fluid was released within 10 msec of a break in the photobeam as the rat licked. Four stainless steel boxes (5 cm X 5 cm X 5 cm), hereafter referred to as “food wells”, were located at the four corners of the experimental chamber and opened to its inside. During the “Lick phase” of the experimental paradigm, the opening to the food wells were blocked by an opaque cardboard panel. During the “Food phase” of the experimental paradigm, the opaque panels were removed and food wells were filled with solid foods available to the rat. Infrared photobeams across the openings to the food wells allowed recording of the time that the rat’s head entered the well.

### Taste and food stimuli

Liquid taste stimuli were chosen to represent the five prototypical taste qualities: 0.1M sucrose (S) for sweet, 0.1M NaCl (N) for salty, 0.0167M citric acid (CA) for sour, 0.0001M quinine HCl (Q) for bitter, and 0.1M monosodium glutamate (MSG) plus inosine monophosphate (0.01M) for umami. All tastants were reagent-grade chemicals (Fisher Scientific, Pittsburgh) dissolved in artificial saliva (AS; 0.015M NaCl, 0.022M KCl, 0.003M CaCl_2_ and 0.0006M MgCl_2_ at a pH of 5.8 ± 0.2; Hirata et al., 2005). AS was also used as a taste stimulus.

Solid foods were also chosen to roughly represent sweet, salty, sour and bitter tastes.

They were: Nestle milk chocolate chips (35% cacao, 67% sugar; sweet), dry roasted salted nuts (salty), Granny Smith apples (sour), and broccoli (bitter). In some animals, other foods were tested: Cheerios (0.036% sugar per serving),), bananas (sweet), Lindt dark chocolate (90% cacao and 0.075% sugar; bitter), shiitake mushrooms (umami) and cheddar cheese (fat).

### Electrode implantation surgery

Approximately 15-20 min prior to induction of anesthesia, subjects were administered buprenorphine (0.01 mg/kg, s.c.) and atropine (0.05 mg/kg, s.c.). Each rat was then placed in an induction chamber and administered isoflurane (3%) for 10 min. Thereafter, the level of anesthesia was maintained with 2% isoflurane throughout the surgery and was adjusted as needed. The scalp was shaved and the head, oriented downward at 25°, was mounted in a stereotaxic instrument (David Kopf Instruments, Model 1900, Tujunga, CA) with blunt ear bars. Artificial tear gel was applied to the eyes to prevent drying during surgery and re-applied as needed. A rectal thermometer attached to a temperature regulator and heating-pad maintained body temperature at 37°C. The scalp was wiped with Betadine alternated with ethanol three times. Once dry, a midline incision was made in the scalp and the fascia were retracted by blunt dissection. Six stainless steel skull screws were mounted in the skull. A hole was drilled between 13.8-15.4mm posterior to bregma and 1.6-2.0mm lateral to lambda. The exposed dura was resected and a 16-channel drivable microwire assembly (23 µm tungsten insulated with polyimide; Innovative Neurophysiology, Durham, NC) was lowered approximately 5.5 mm ventral to the brain’s surface and cemented in place with dental acrylic. A stainless steel wire was wrapped around a skull screw to serve as ground. The implanted electrode assembly and skull screws were embedded in dental acrylic. Anesthesia was then discontinued, and the rat was placed in its home cage until it was conscious and mobile. Thereafter, 0.05 mg of buprenorphine, 3 mL of warmed saline, and 0.05 mg gentamicin were administered s.c. A topical antibiotic (Neosporin) was also spread on the skin around the headcap. Buprenorphine and gentamicin were given daily for three days post-surgery. Rats were weighed daily until they had returned to 90% of their pre-surgical body weight.

### Experimental procedures

*Overview* Once recovered from electrical implantation surgery, rats were water deprived for 18-23 hrs and trained to lick sucrose from the lick spout in the experimental chamber. This training consisted of daily (except for weekends) 30 min sessions of presentation of 0.1 M sucrose dissolved in AS. Once the rat achieved > ∼500 licks/d for three days, electrophysiological recording sessions were begun. The water deprivation regimen was continued throughout the recording sessions.

Each experimental session consisted of two paradigms: the Lick paradigm and the Eating paradigm, as described below. Some animals did not receive the Lick paradigm, but when it when administered, the Lick paradigm always preceded the Eating paradigm. Each paradigm was 30 min long and separated by 1hr or less. A Cineplex Behavioral Research system (Plexon, Inc., Dallas, Texas) synchronized electrophysiological and video recordings of all behaviors during both experimental paradigms. Neural data and timestamps of stimulus events were acquired through an Omniplex server system (Plexon, Inc., Dallas, Texas).

*Lick paradigm.* Rats were connected to the Omniplex system through a headstage and cable connected to a commutator and placed in the operant chamber for 30 min. Rats were given access to a sipper tube where tastants were dispensed. Tastant trials consisted of five consecutive licks of a taste stimulus, followed and preceded by six AS licks presented on a variable ratio 5 schedule, with dry licks interspersed between AS licks. The order of presentation for the taste stimuli was randomized.

*Eating paradigm*. Following the Lick paradigm, rats were either left in the experimental chamber or removed to a holding cage while the food wells were loaded with 3 gms of solid food. The positions of the various foods were pseudo-randomized across days. Some sessions included an empty well. All foods were chopped and weighed daily before placement into wells. Rats were given 30 min access to food. When the session was completed, the remaining food was weighed and recorded.

### Data Analysis

#### Spike sorting and unit identification

Neuronal waveforms were isolated through in-house Matlab software (The MathWorks, MA) or by OfflineSorter (Plexon, Inc., Dallas, Texas). Neurons were considered isolated based on the following criteria: An acceptable L-ratio (−*log*_10_(*Lratio*) ≥ 1), signal to noise ratio (SNR ≥ 3), and percentage of interspike intervals ≤ 1 ms (ISI ≤ 0.5%). (Schmitzer-Torbert, Jackson, Henze, Harris, & Redish, 2005). The L ratio is a measure of how well spikes in a cluster are separated from other spikes recorded from the same electrode.

To verify that recorded neurons were the same between the Lick and Eating paradigms, we compared waveform templates. For the recording from each paradigm, we averaged waveform matrices (n x 32 dimensions) across rows, forming a row vector with a dimensionality of 32. Both waveform templates belonging to the Lick and Eating paradigms were then overlaid on one another and a Pearson’s correlation coefficient was computed. To be classified as the same unit in both experimental paradigms, a Pearson correlation coefficient between waveform templates ≥ 0.99 was required (see Sammons et al., 2016).

### Lick paradigm - taste response detection and classification

In the Lick paradigm, taste responses were classified as either lick by lick (LxL) or five lick (5L) as described previously (Roussin et al. 2012). LxL responses were short in duration (≤150 ms) and occurred between individual licks. 5L responses were generally longer in duration and spanned two or more licks. A Chi square test was used to determine whether a taste response was LxL. The Chi square test used the peri-stimulus time histogram (PSTH, 15 ms bins) of the post stimulus period for tastant licks compared with the PSTH for dry licks (non-reinforced). The post-stimulus period (150 msec) for each tastant lick was the predictor and the post stimulus period for dry licks was the expected. If the predictor was significantly different from the expected (*p* < 0.05), a response was classified as a LxL response.

For 5L response detection, responses were measured by comparing the post-stimulus period (4 s) for each tastant trial to the pre-stimulus period (baseline; 2 s prior to the first tastant lick). A 95% confidence interval was calculated based on the average baseline firing rate. For each post-stimulus tastant trial, spikes were binned into row vectors with a dimensionality of 200 (each element was a 20 msec bins). A sliding window of length 100 msec incremented by 20 msec along the 200-dimensional vector. If three consecutive bins were above or below the 95% confidence interval, then the trial was classified as significant. At least half the trials for each stimulus needed to be significantly above or below the baseline confidence interval for a stimulus response to be considered significant. This response detection methodology was not used on tastants with less than four trials.

### Eating paradigm – behavioral event and neural response detection

*Scoring of eating related events:* The timing of eating and grooming events was scored offline by one or more raters/observers. Inter-rater reliability was assessed by analyzing a subset of sessions that had multiple raters. For each behavioral event that was identified, and for each rater, we computed the start time of the interval during which the event occurred. We then computed the average ± SEM difference of the start times scored for each event across raters. If one rater scored an interval and another had not, that interval was removed from the inter-rater calculation. For example, for the session shown in Fig. 2, the average start time difference between scored eating events was 100 ms ± 121 ms.

**Figure 1.**
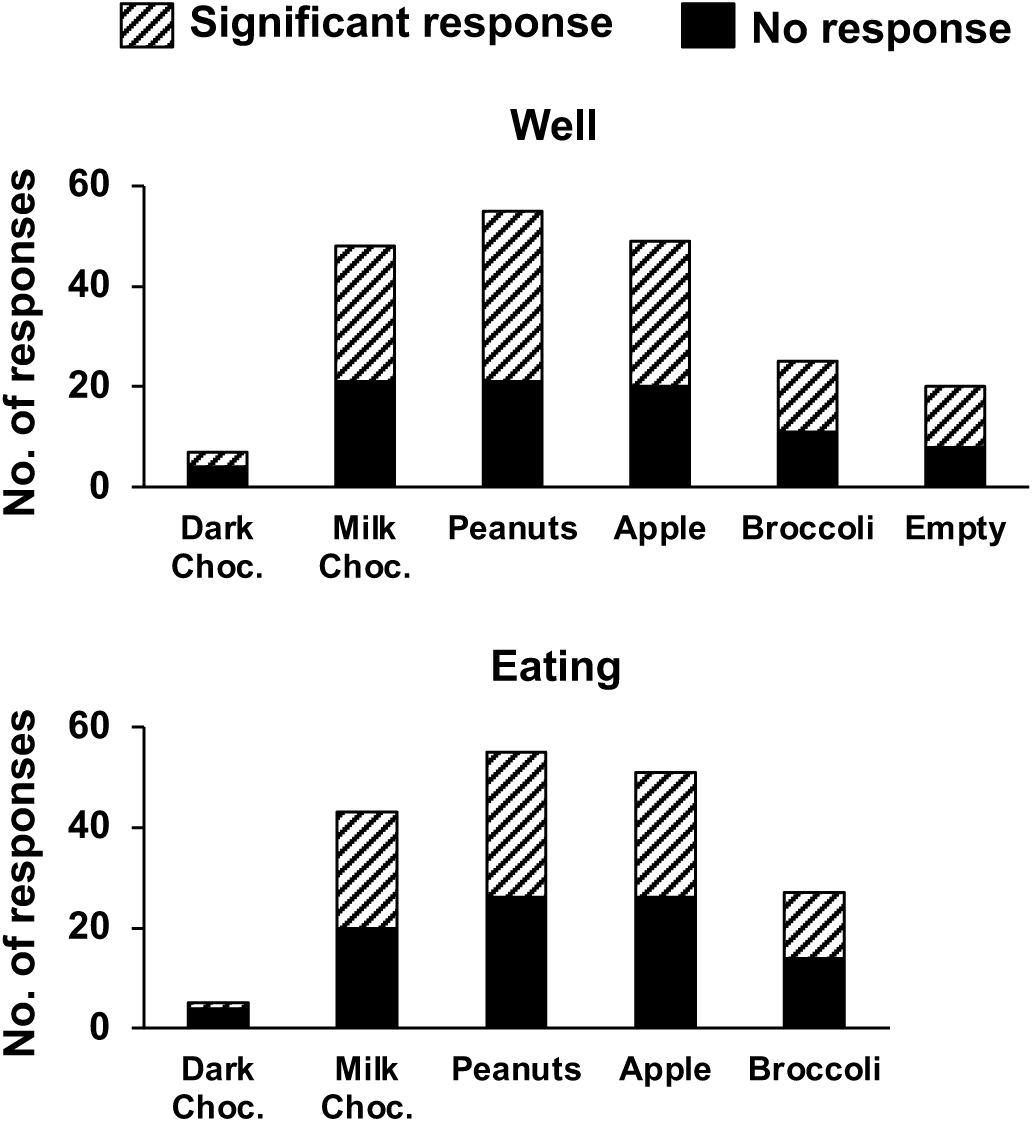
Number of rNTS responses to food well exploration (top) and eating. For each food, and the empty well, the proportion of significant responses is also indicated. Not all foods were tested in any one given session. See text for details.

**Figure 2.**
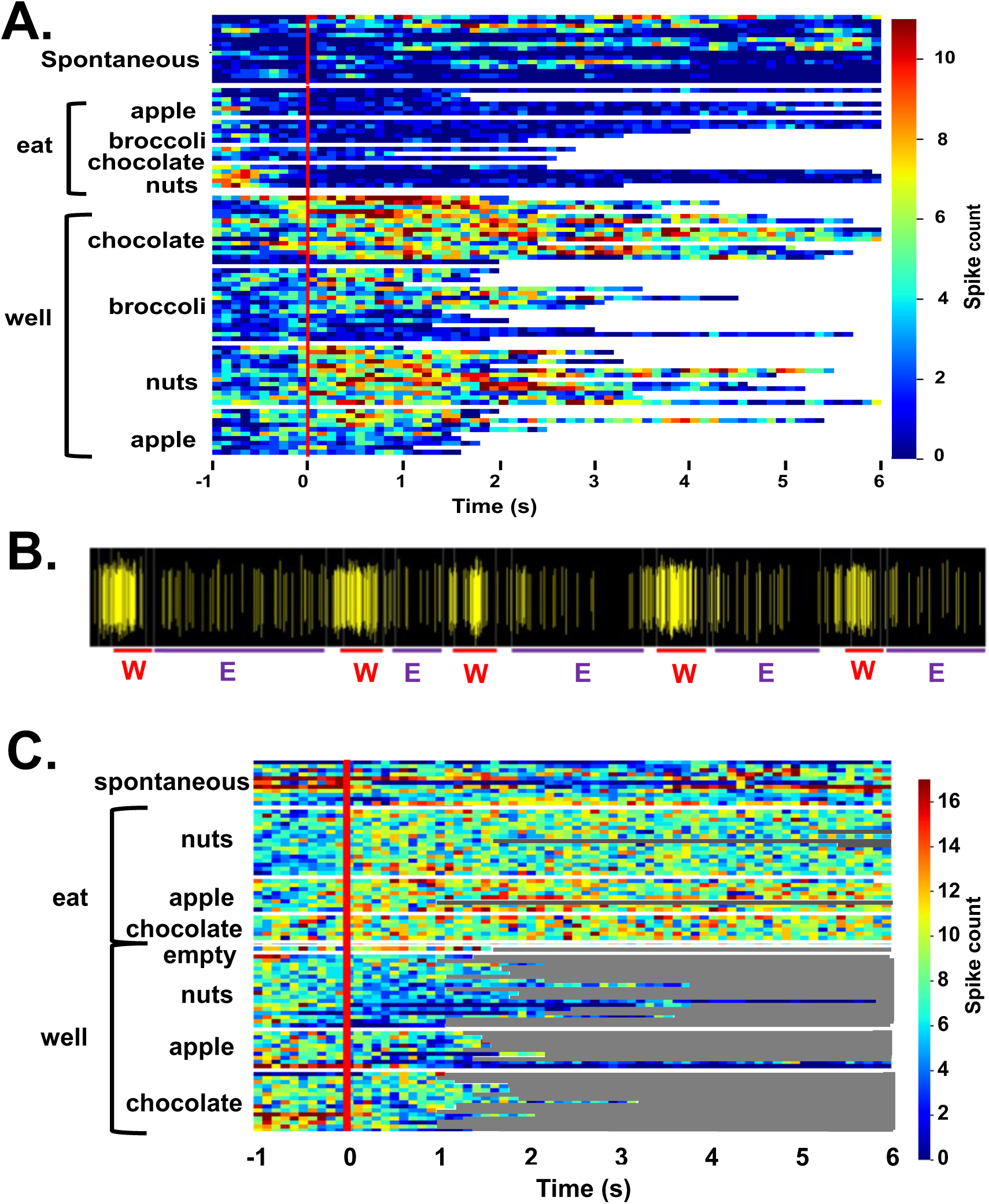
A. Heat map showing the response of a single unit in the rNTS to exploration of the food-filled web and to eating. Spontaneous firing rate, preceding the onset of each event (vertical red line) is also shown. Each line represents one trial and its length represents the time that the animals engages in the respective activity. This cell becomes active just prior to head entry into the well and is nearly silent during eating. B. Spiking activity of another cell during well exploration (W) and eating (E). The cell begins to fire vigorously just prior to the animal’s head breaching the threshold of the well, prior to any contact with the food. C. Heat map showing the activity in a cell that is excited when the animal is eating, but not when the animal is exploring food in the well.

*Solid food response detection*: Significant responses to solid food were determined by comparing the firing rate during the post-stimulus period (1 s; well exploration or food consumption) to the spontaneous firing rate. Spontaneous activity was defined as activity that was at least 2.5 seconds away from well entry and eating events to avoid any solid food induced residual activity. Intervals of spontaneous activity varied in length across the eating paradigm. Since spontaneous firing rates were so variable across the session, responses to well exploration or food consumption were considered significant if the average firing rate was either above or below the spontaneous rate by at least five spikes per second (sps). For some analyses, the spontaneous firing rate was subtracted from the response firing rate, as noted below.

*Change point analysis.* Change point analysis of firing rate was used to assess the onset of responses just prior to well entry. Specifically, we considered the spiking activity from 2 s prior to well entry to 4 s following well entry in 100 ms bins. The change point detection was that described by Arif et al. (2017). If a single change point was detected prior to well entry, we considered this to be a response.

*Multidimensional scaling (MDS) analyses.* To assess the inter-relationships of the across unit patterns of response for well entry and exploration, and eating, an MDS analysis was performed on the average evoked firing rates for each food across trials for each cell using XLSTAT. The similarity matrix was formed using Pearson *r* correlations and the Kruskal stress test was used to determine the number of dimensions that best fit the data. Average firing rates across trials evoked by well entry and eating episodes minus spontaneous firing rate were included for each cell and for each food and the empty well whether or not they varied by more than five sps from spontaneous firing rate.

### Other Statistical analyses

For assessing differences in spontaneous firing rate between sessions, a paired t-test was used. A standard linear regression was also used for assessing the relation between lick and solid food spontaneous firing rates. Difference between solid-food response magnitudes were analyzed with a Kruskal-Wallis test. A rank-sum test assessed difference between well-entry and eating entropy.

## Results

Electrophysiological responses to the approach, sampling and eating of various solid foods were recorded in 60 NTS cells from eight awake, unrestrained rats. Fig. 1 shows the number and proportion of responses to each of the foods tested. Of these 60 cells, 28 (47%) were also recorded in a lick paradigm for responses to traditional taste stimuli preceding the eating paradigm. Although 55 taste-responsive cells were recorded overall in the lick paradigm, only about half remained well isolated throughout the eating phase of the experiment. A total of 50 sessions were given, with animals given 6.7±1,4 sessions on average (range = 2-14). All but three Eating sessions were preceded by Lick sessions.

### Behavioral observations

When rats were given the opportunity to approach, sample and eat various solid foods, the pattern of behavior was consistent and stereotypical. Rats approached the food well with their heads leaning into the well. Once the head entered the well, there was lick-like mouthing accompanied by grabbing of small bits of food that were eaten while the head was in the well.

Larger morsels were grabbed with either the mouth or paws and eaten outside of the well, either just outside the well or toward the center of the experimental chamber. These behavioral patterns were idiosyncratic. Some rats roamed, apparently randomly, from food well to food well, eating the various foods along the way. Others stayed at one food well and ate the contents until the well was empty and then moved on to the next food well. In most cases, rats poked their head and forepaws into a well briefly and sampled and/or grabbed food which they ate outside of the well. Data presented in Table 1 show that, on average, rats spent about 3 s in the well and between 7.5 and 20 s eating food outside of the well, depending on the food. When peanuts, milk chocolate, apples and broccoli were presented simultaneously, there were no differences in either the number of visits to the food wells across foods (ANOVA, *F*(3, 76) = 1.810; *p* = 0.1525) or the time spent in the wells (ANOVA, *F*(3, 76) = 2.212; *p* = 0.936). Similarly, there were no differences among foods in the number of eating events when these foods were presented simultaneously (ANOVA, *F*(3, 76) = 1.858; *p* = 0.1438). However, there was a significant difference in the time spent eating these foods (ANOVA, *F*(3, 76) = 2.855; *p* = 0.0427), with less time spent eating milk chocolate than apple (pairwise comparisons via Tukey’s multiple comparisons test, *p* = 0.0306). For 20 sessions, an empty well was presented along with three wells that contained food. Although the rats visited the empty well a few times in these sessions, the dwell time was generally less than that for food-filled wells. However, in one outlier session the rat visited the empty well 36 times and explored it extensively (mean = 6.3 ± 0.03 s).

**Table 1.** Trial numbers and dwell times.

**A. Well. No. of trials (Mean ± SEM) and Dwell time (s) (Mean ± SEM)**
|  | Dark choc. | Milk Choc. | Peanuts | Apple | Broccoli | Empty |
| --- | --- | --- | --- | --- | --- | --- |
| No. trials | $6.4 \pm 1.3$ | $15.0 \pm 1.0$ | $14.0 \pm 1.1$ | $11.2 \pm 1.2$ | $9.3 \pm 0.8$ | $5.2 \pm 1.9$ |
| Dwell time | $3.2 \pm 0.3$ | $2.5 \pm 0.2$ | $2.9 \pm 0.1$ | $3.0 \pm 0.6$ | $2.5 \pm 0.1$ | $2.3 \pm 0.5$ |
| No. of cells | 11 | 60 | 56 | 53 | 32 | 19 |

**B. Eating. No. of trials (Mean ± SEM) and Dwell time (s) (Mean ± SEM)**
|  | Dark choc. | Milk choc. | Peanuts | Apple | Broccoli |
| --- | --- | --- | --- | --- | --- |
| No. trials | $4.8 \pm 1.1$ | $9.8 \pm 0.8$ | $11.2 \pm 1.3$ | $8.9 \pm 0.7$ | $6.6 \pm 0.9$ |
| Dwell time | $20.3 \pm 6.4$ | $8.2 \pm 0.6$ | $14.3 \pm 1.4$ | $15.0 \pm 1.6$ | $9.2 \pm 1.2$ |
| No. of cells | 9 | 57 | 51 | 50 | 28 |

\*Some entries to the food wells were not counted because they were $< 1$ s.

### Electrophysiological responses to approach, sampling and eating food

Approach and time spent in the food wells generally resulted in vigorous and robust increases in firing rate that varied across food types. In contrast, many cells showed significantly attenuated firing rates when the animal was eating.

Fig. 2 shows examples of responses to well exploration and eating in three cells. The cells whose activity is shown in Fig. 2A and 2B are excited when the rat enters the well, but not when it consumes the food. The cell in Fig. 2B increases its firing rate just prior to the moment when the rat’s head breaches the threshold of the well, before any contact with the food. The cell in Fig. 2C shows an excitatory response to eating the food, but shows little response to well exploration. Table 2A summarizes the responses to well entry and or eating. For milk chocolate, peanuts, and apple, there were twice as many cells that responded exclusively to well entry than those that responded only when the animal was consuming those foods. Table 2B-E summarizes the electrophysiological responses across well entry and eating episodes." As can be seen, excitatory responses to well entry were more prevalent than inhibitory responses but the opposite was true for eating episodes. Overall, average firing rates in response to both well entry and eating were generally equivalent across foods.

**Table 2.**
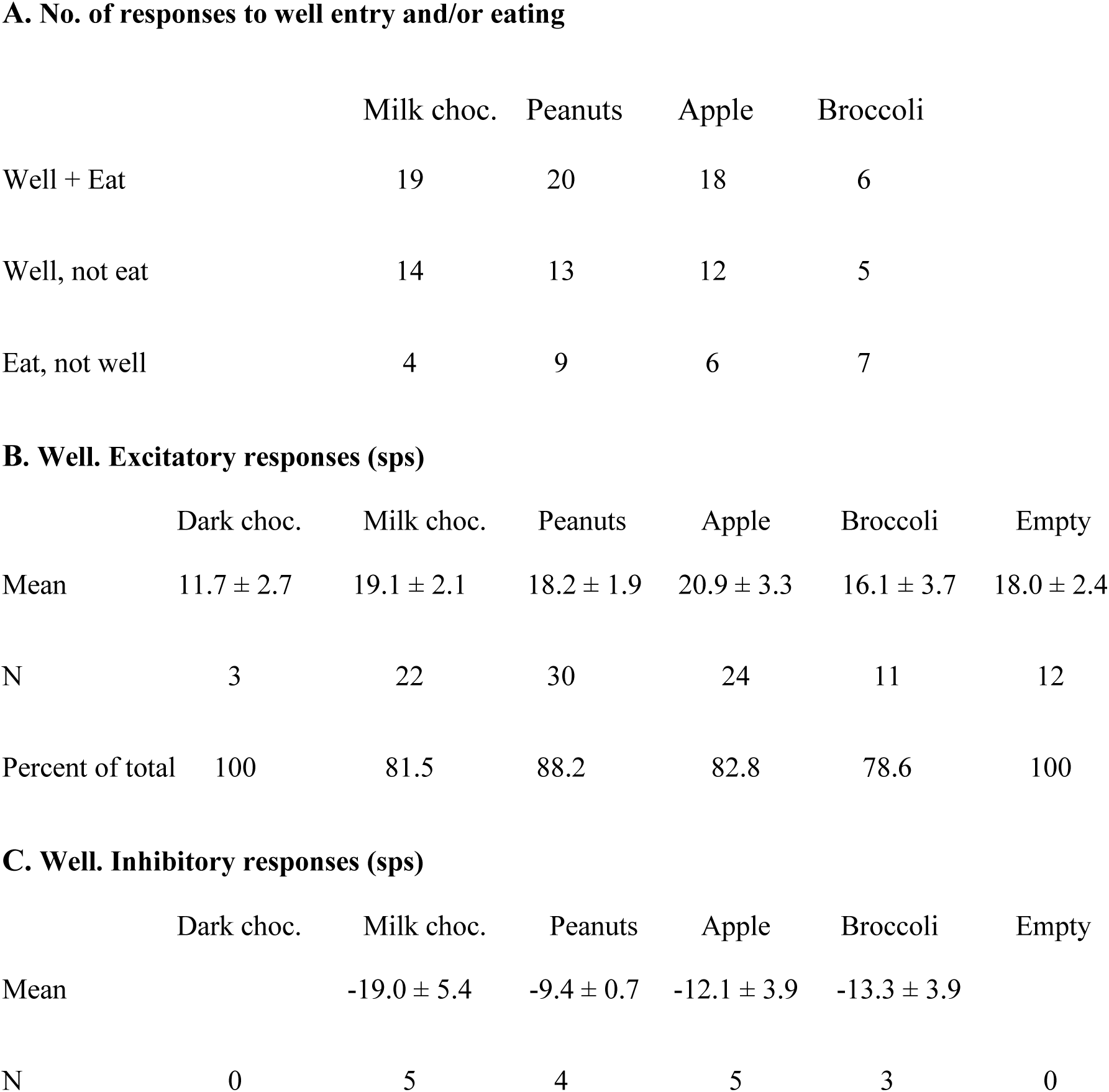
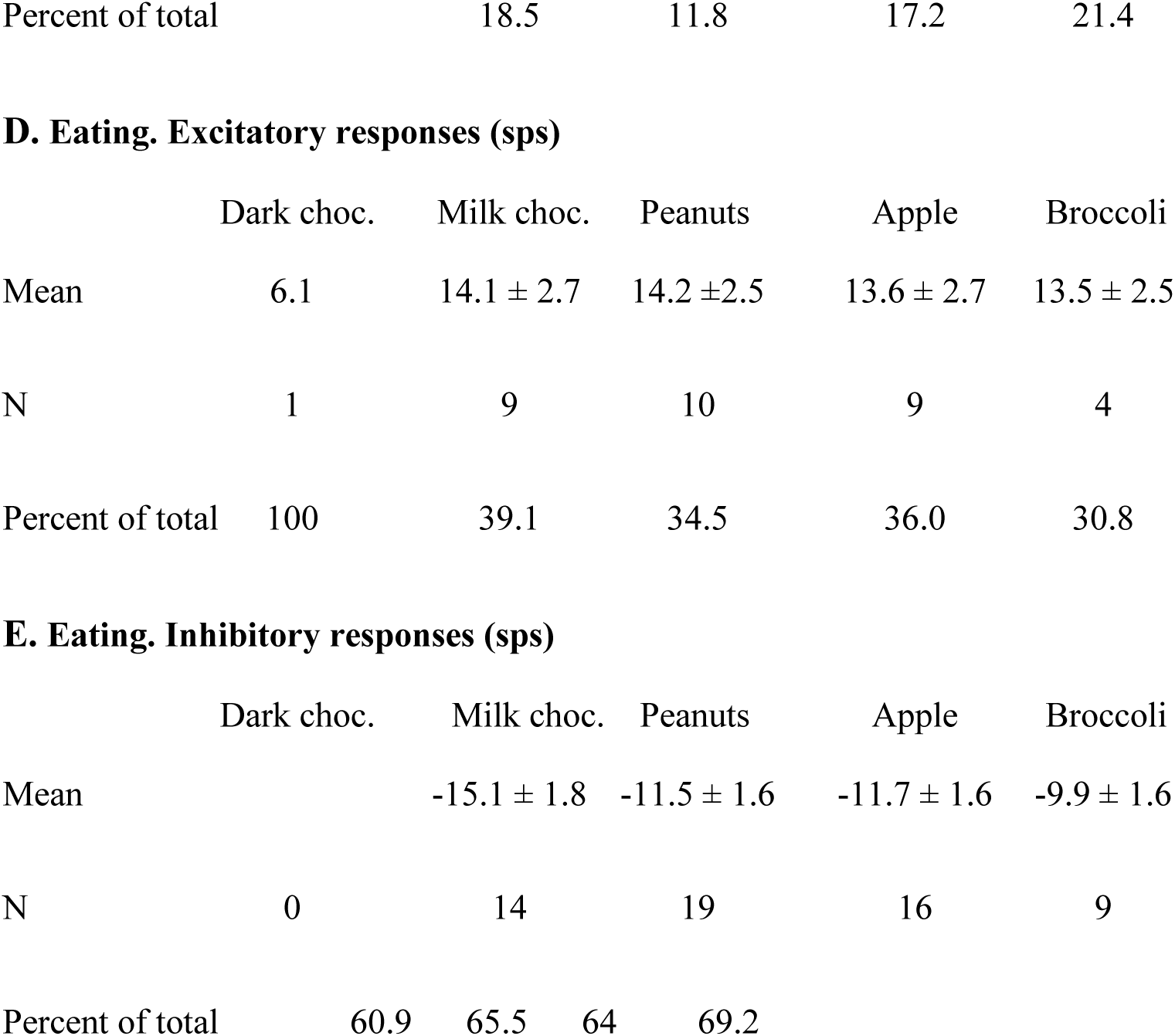
Electrophysiological responses to food well and eating.

Fig. 3 summarizes the firing-rate responses to well entry and eating in individual cells for the most frequently tested foods: salted peanuts, milk chocolate, Granny Smith apples and broccoli. Firing rates while eating were almost universally lower than those during well entry, even though not all eating responses were inhibitory. Specifically, for peanuts, 80% (16/20 responses to both well and eating), milk chocolate, 80% (16/20 responses), for apple, 61% (11/18 responses) and for broccoli 67% (4/6 responses) of the responses to eating were reduced by more than 5 sps below responses to well entry. Analyses of response magnitudes evoked by well entry vs. eating in cells that were tested in both conditions to the same food (spontaneous firing rate included) showed that, for the four foods that were tested most frequently, responses to well entry were significantly larger than that to eating. Specifically: for peanuts, *t* = 5.048, *df* = 41, *p* < 0.0001; for milk chocolate, *t* = 5.844, *df* = 51, *p* < 0.0001; for apples, *t* = 4.587, *df* = 47, *p* < 0.0001; for broccoli, *t* = 3.832, *df* = 23, *p* < 0.0009. For dark chocolate, *t* = 1.982, *df* = 4, *p* = 0.1186. Moreover, data shown in Table 3A shows that about half of the sample of cells responded to well entry or eating but not both, implying that there may be a subset of cells that are specialized to respond to the appetitive or consummatory aspects of food.

**Figure 3.**
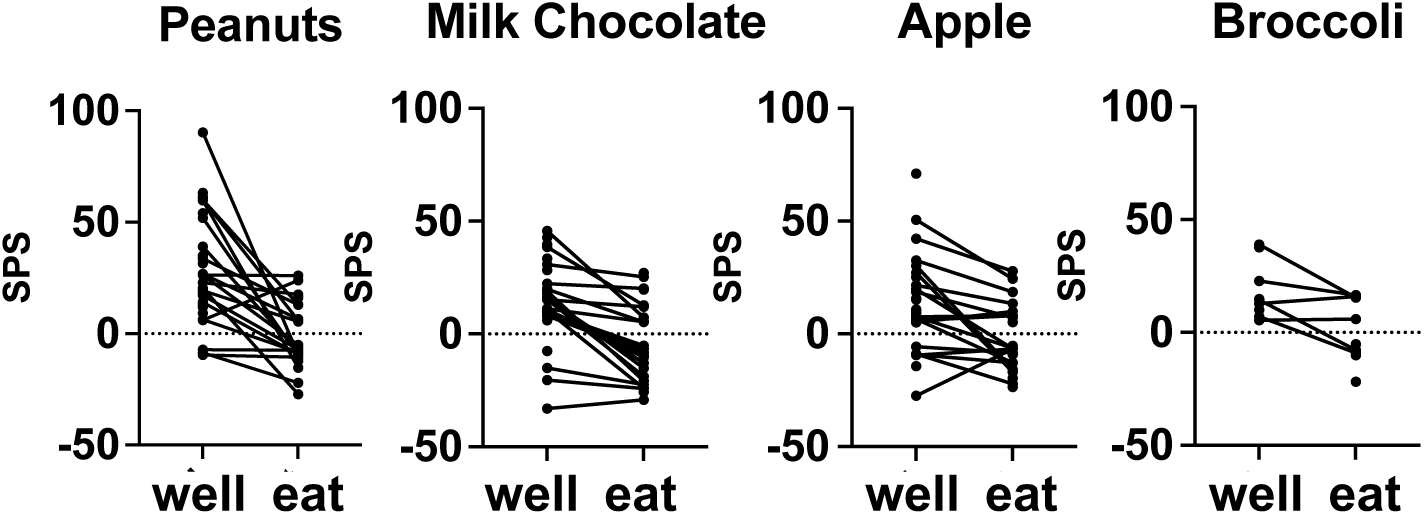
Mean firing rate (sps) evoked by well exploration and eating in cells that were tested with salted peanuts, milk chocolate, Granny Smith apples, and broccoli. Data from cells that were tested with peanuts, chocolate, and apples, but not broccoli, were also included. Most cells reduced their firing rate when the animal was eating, compared to when the animal was exploring the food-filled well. See Table 2 and text for details.

**Table 3.**
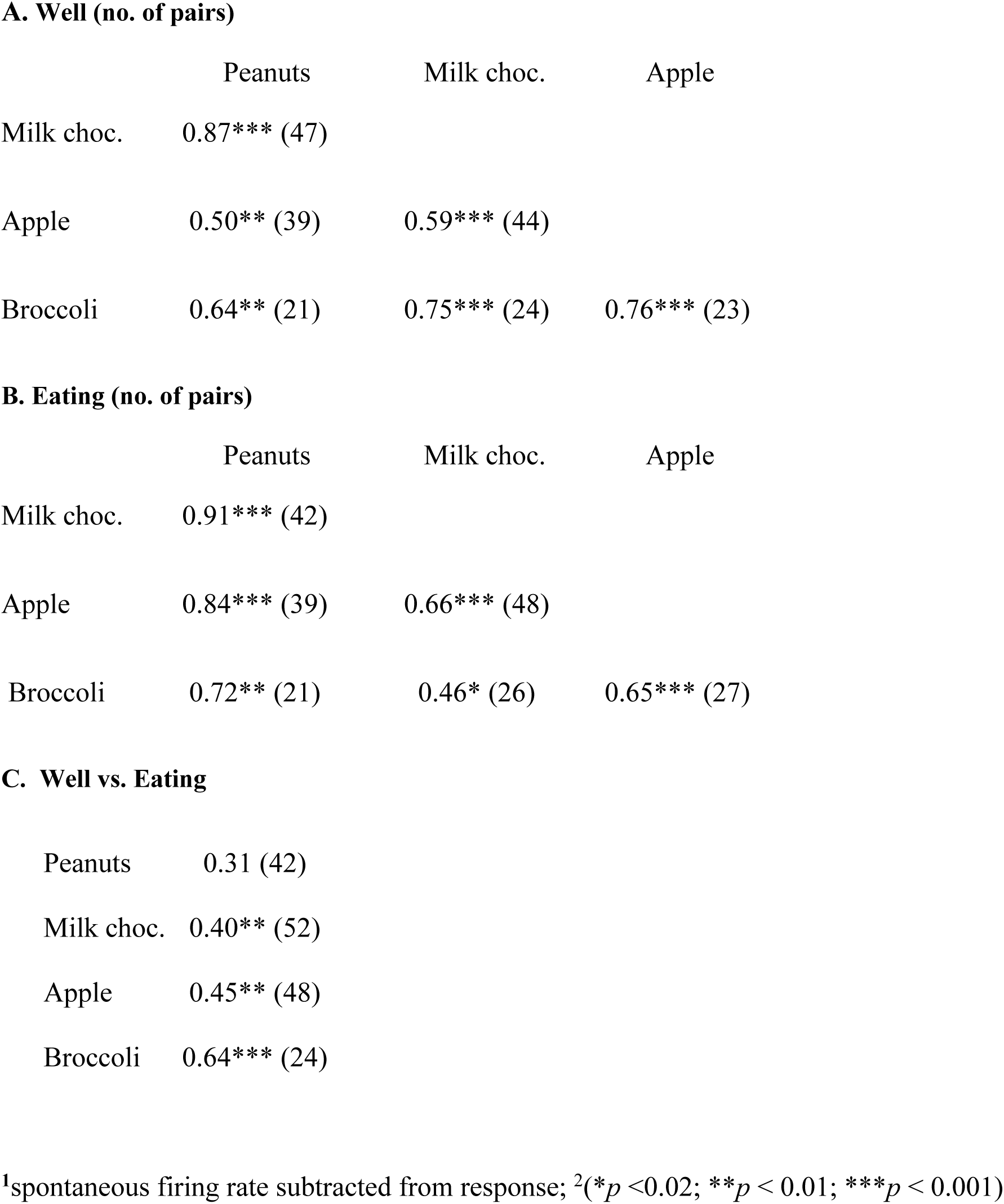
Well vs. Eating response correlations^1,2^.

**A. Well (no. of pairs)**
|  | Peanuts | Milk choc. | Apple |
| --- | --- | --- | --- |
| Milk choc. | 0.87*** (47) |  |  |
| Apple | 0.50** (39) | 0.59*** (44) |  |
| Broccoli | 0.64** (21) | 0.75*** (24) | 0.76*** (23) |

**B. Eating (no. of pairs)**
|  | Peanuts | Milk choc. | Apple |
| --- | --- | --- | --- |
| Milk choc. | 0.91*** (42) |  |  |
| Apple | 0.84*** (39) | 0.66*** (48) |  |
| Broccoli | 0.72** (21) | 0.46* (26) | 0.65*** (27) |

**C. Well vs. Eating**
|  |  |
| --- | --- |
| Peanuts | 0.31 (42) |
| Milk choc. | 0.40** (52) |
| Apple | 0.45** (48) |
| Broccoli | 0.64*** (24) |

<sup>1</sup>spontaneous firing rate subtracted from response; <sup>2</sup>(\* $p < 0.02$ ; \*\* $p < 0.01$ ; \*\*\* $p < 0.001$ )

Responses within a cell to both well entry and eating most often varied according to the food being explored (well) and/or eaten (eat), and the foods that elicited the largest response – either at well entry or during eating – were broadly distributed. With regard to well entry, among the 20 cells tested with four foods, there were five cells that responded best to peanuts, seven cells to milk chocolate, four cells to apple and four cells to broccoli. During eating episodes, there were three cells that responded best to peanuts, five to milk chocolate, five to apple and seven to broccoli. However, only three cells showed the same best stimulus in both well exploration and eating; the best stimuli for these foods were nuts, milk chocolate and broccoli. Similar results were found for the 17 cells that were tested with only three foods.

Fig. 4 summarizes the tuning of all cells tested with either three or four foods as described above, subdividing the population according to the best food stimulus, considering separately the well-entry phase (left) and the eating phase (right). Each response was calculated as a percentage the response that evoked the largest response in that phase. It is apparent that the majority of cells did not respond equally to all foods tested – and in many cases, cells that responded with an increase in firing rate to one food responded with a decrease in firing rate to another.

**Figure 4.**
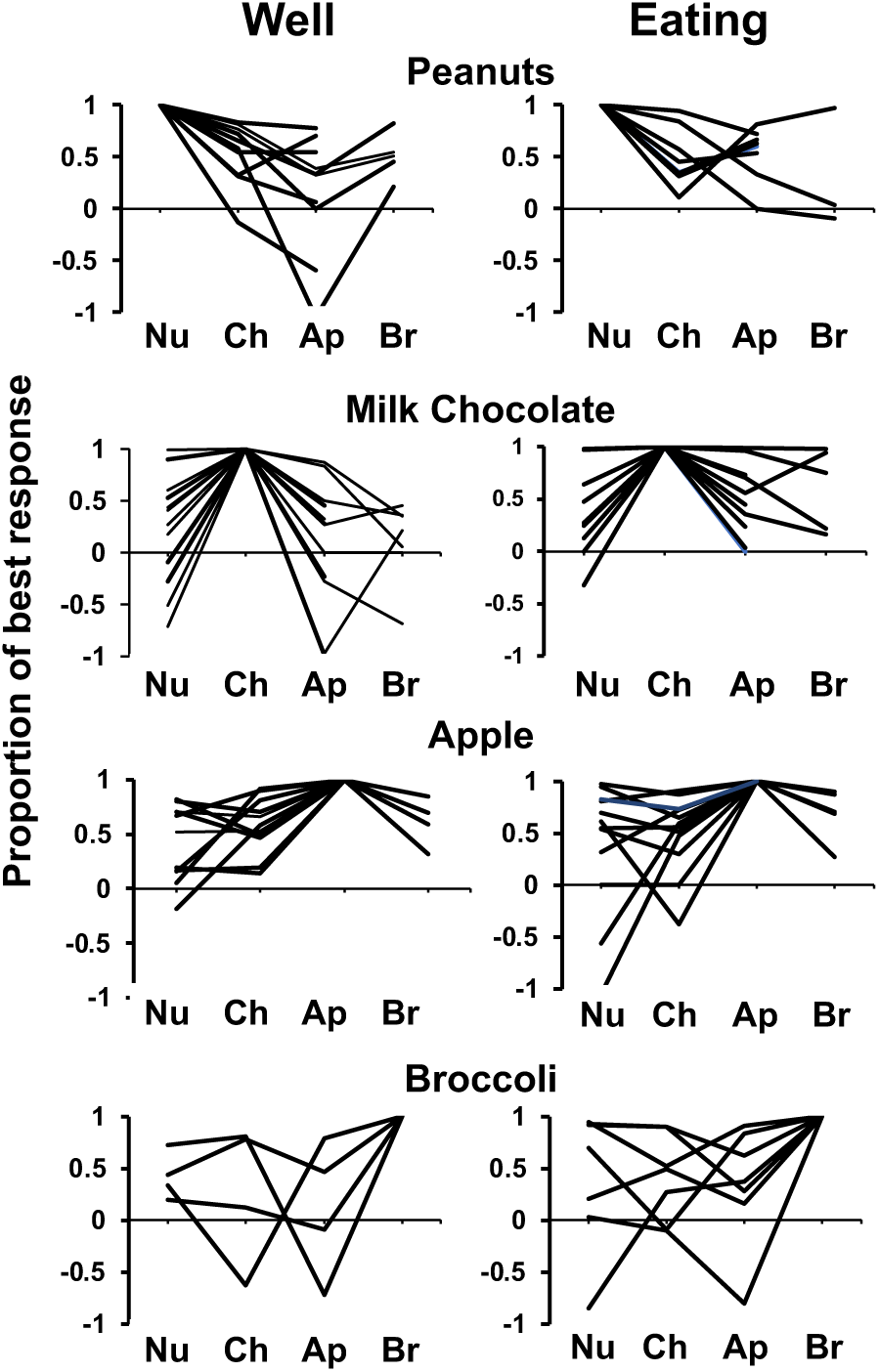
Response profiles of cells that responded best to each salted peanuts, milk, chocolate, Granny Smith apples, and broccoli and each condition (well exploration and eating). Responses are expressed as a proportion of the most vigorous response, with negative values indicating firing rates below baseline. Cells that were tested with all four foods, as well as those tested without broccoli are included. Most cells show some degree of specificity. Abbreviations are as follows: Nu, salted peanuts; Ch, milk chocolate; Ap, Granny Smith apples; Br, broccoli.

On the other hand, many cells were broadly tuned across foods. We quantified the breadth of tuning using the standard Sharpness Index (Rainer et al. 1998) as follows:

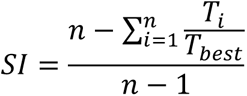

where *SI =* Sharpness Index, *n* = number of stimuli, and *T* = the mean response magnitude for a given stimulus. A value of 1.0 signifies narrow tuning while value of 0 indicates broad tuning across stimuli. Note that, because many responses that were below spontaneous firing rates, the Sharpness Index was conducted using raw response magnitudes, rather than after subtraction of spontaneous firing rates. This necessarily pushed Sharpness Index measures downward, perhaps artificially imposing a bias toward more broad tuning. Nevertheless, relative selectivity measures were informative. For responses in cells that were tested with either three or four stimuli (*n* = 38, see Fig. 5), the Sharpness Index for well entry was on average 0.21 ± 0.02, for eating 0.27 ± 0.03. Though there was no statistically reliable difference between the Sharpness Indices for well entry compared with that for eating (paired *t* test, *p* = 0.0697), 37% (15/38) of cells showed narrower tuning, i.e., an increased Sharpness Index by at least 0.1), during eating vs. well entry. In contrast, only 13% (5/38) became more broadly tuned during eating.

**Figure 5.**
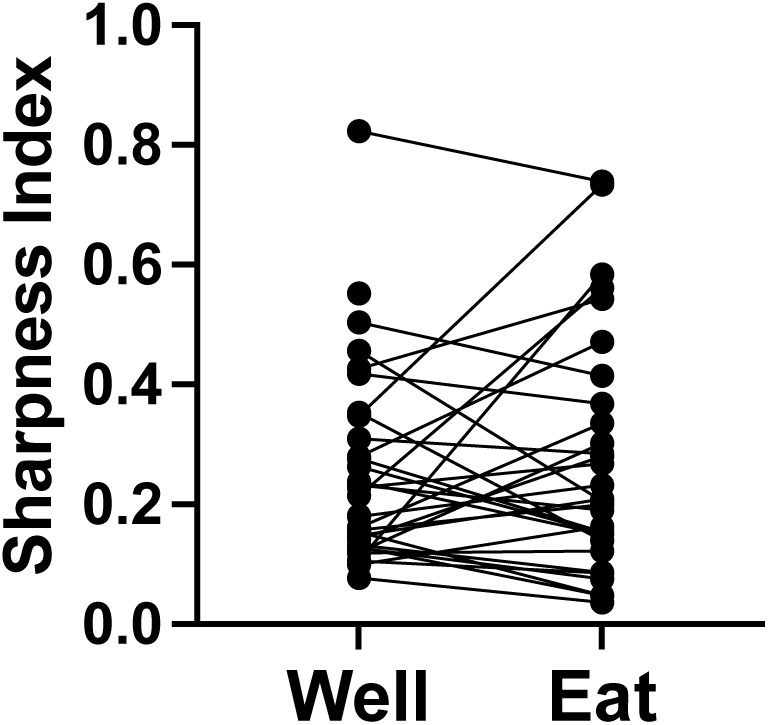
Sharpness Index (a measure of tuning) for cells tested with four (salted peanuts, milk chocolate, Granny Smith apples, and broccoli) or three (peanuts, chocolate, and apples) foods. (*N*=38)

Analysis of across-unit patterns of rNTS responses (Table 3) shows substantial commonalities of neural tuning for both well and eating events. In addition, responses to well events were significantly positively correlated with responses to eating events for milk chocolate, apple, and broccoli, but not for peanuts. In general, these data suggest that the across unit pattern of responses to these foods does not discriminate well among food types, either in the appetitive of consummatory phase, and that the cells that respond to both well entry and eating do so in similar ways.

Results of MDS analysis, shown in Fig. 6 provides additional insights. The across-unit patterns for all foods during the appetitive phase are separated from those during the eating phase, separated in a direction that is approximately perpendicular to the predominant directions within each of these subsets. Within each subset, the across unit patterns in the appetitive phase were more similar to each other than those evoked during the eating phase, and responses to the empty well were segregated from those generated by all foods. These patterns suggest that consummatory phase, and food quality, are independently represented at the population level.

**Figure 6.**
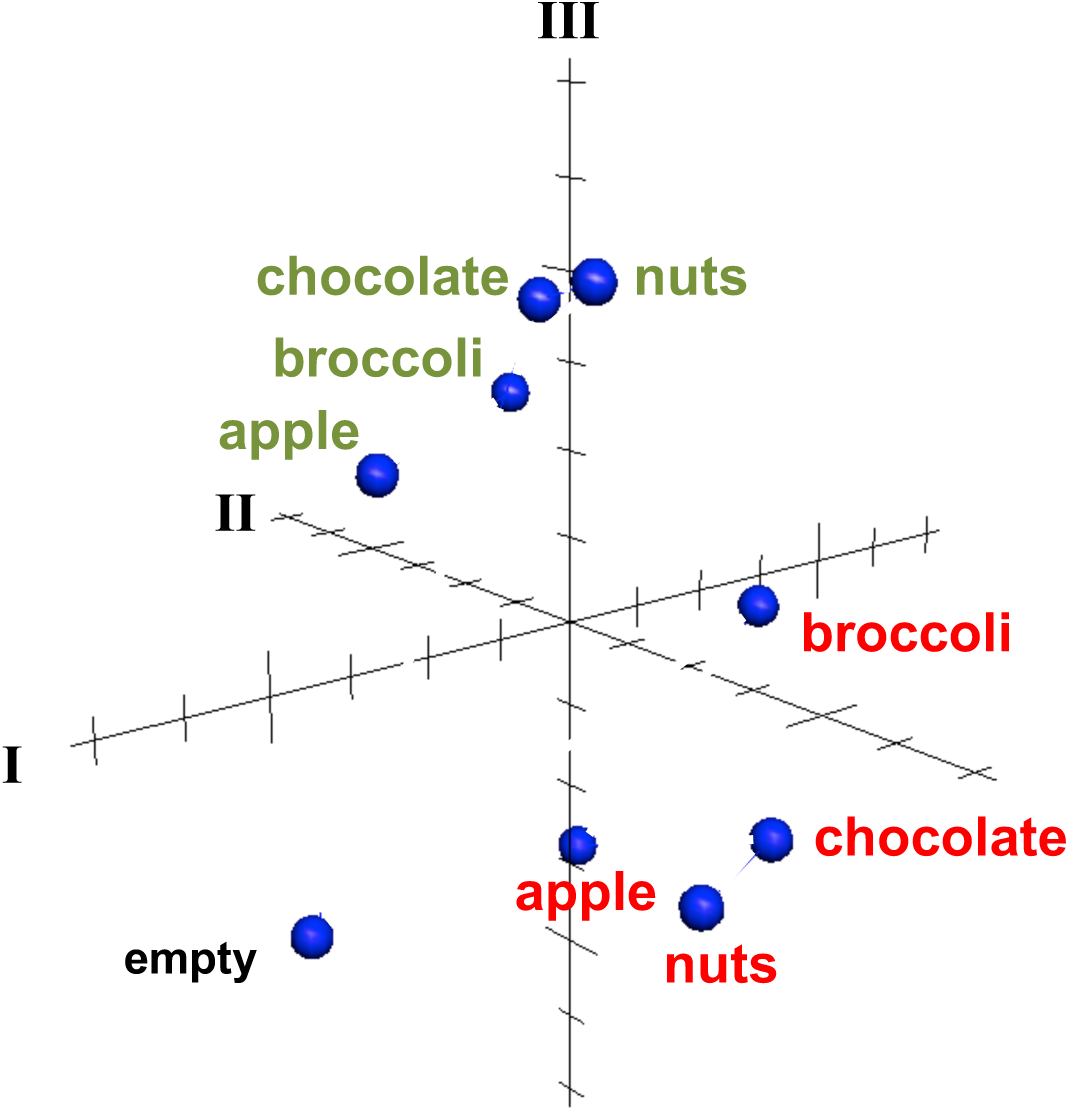
MDS analysis of across unit patterns evoked by food-filled well exploration (red) and eating (green) as well as exploration of an empty well. Response patterns evoked by well exploration are clustered separately from those evoked during eating. The across unit pattern evoked by an empty well is segregated from both eating and well exploration clusters. See text for details.

### Relationship between electrophysiological responses to traditional tastants and food

Forty-seven of the 50 sessions of the Eating paradigm were preceded by a session of the Lick paradigm, and of the 60 cells recorded in the licking paradigm, recordings in 28 cells were still present during the eating paradigm. Twenty-two of these cells were taste responsive, that is, they responded to at least one of the taste stimuli tested. Fig. 7 shows an example of the responses to taste stimuli as well as the responses to well exploration and eating food. This cell responded to MSG and weakly to NaCl and sucrose in the Lick paradigm. When the animal was permitted access to solid food, this cell responded to well exploration for all the foods tested, but did not respond when these foods were eaten (panels B and C). A sample of the spiking activity in this cell shows that it began to respond to well entry prior to any contact with the food (panel C). Fig. 8 (top) shows the distribution of cells responsive to 1-5 of the tastants tested. The majority of cells (67%) responded to more than one of the prototypical taste stimuli representative of the basic taste qualities: sweet, salty, sour, bitter and umami. Fig. 8 (bottom) shows the mean taste response magnitudes for each taste stimulus as well as the response profiles for all of the taste-responsive cells. Six cells were completely unresponsive to any of the taste stimuli tested but nevertheless responded to food. Fig. 9 shows an example of a non-taste-responsive cell, detailing its response to the traditional taste stimuli and its response to food.

**Figure 7.**
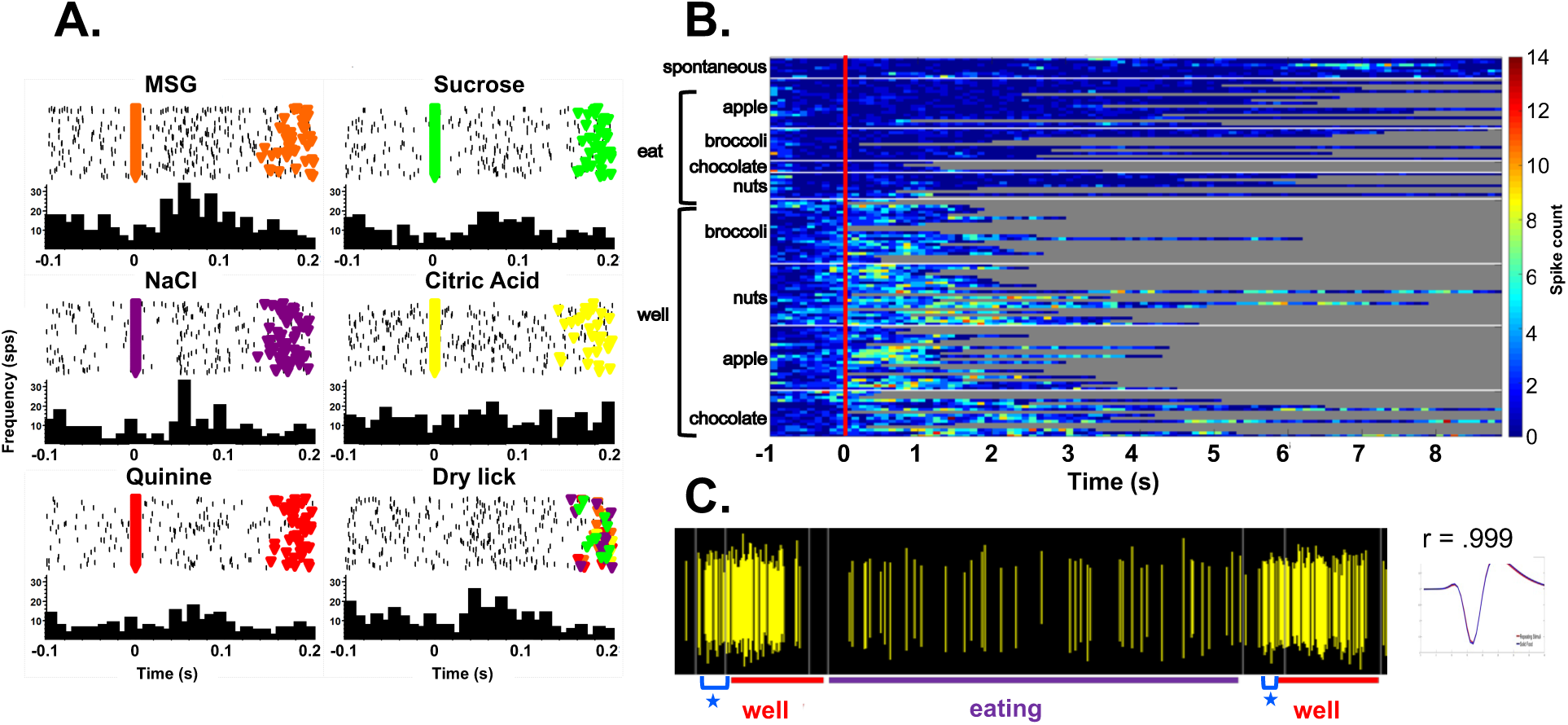
A. Peristimulus-time histograms (PSTHs) responses to tastants in one cell as the animal licked in the Lick paradigm. Top of each panel shows a raster plot where each line represents one trial. Bottom of each panel shows the resulting histogram of unit activity. This cell responded to well to MSG and briefly to NaCl. B. Heat map of responses to well exploration and eating in the cell shown on the left. There was a robust response to well exploration, especially for nuts and apples; however, the cell returned to spontaneous firing rate during eating. C. Spiking activity in the same cell showing an anticipatory response to well entry denoted by a blue star. This anticipatory activity occurred prior to any contact with the food. The waveform evoked by this cell during well exploration and eating are superimposed and shown to the right of the spiking activity.

**Figure 8.**
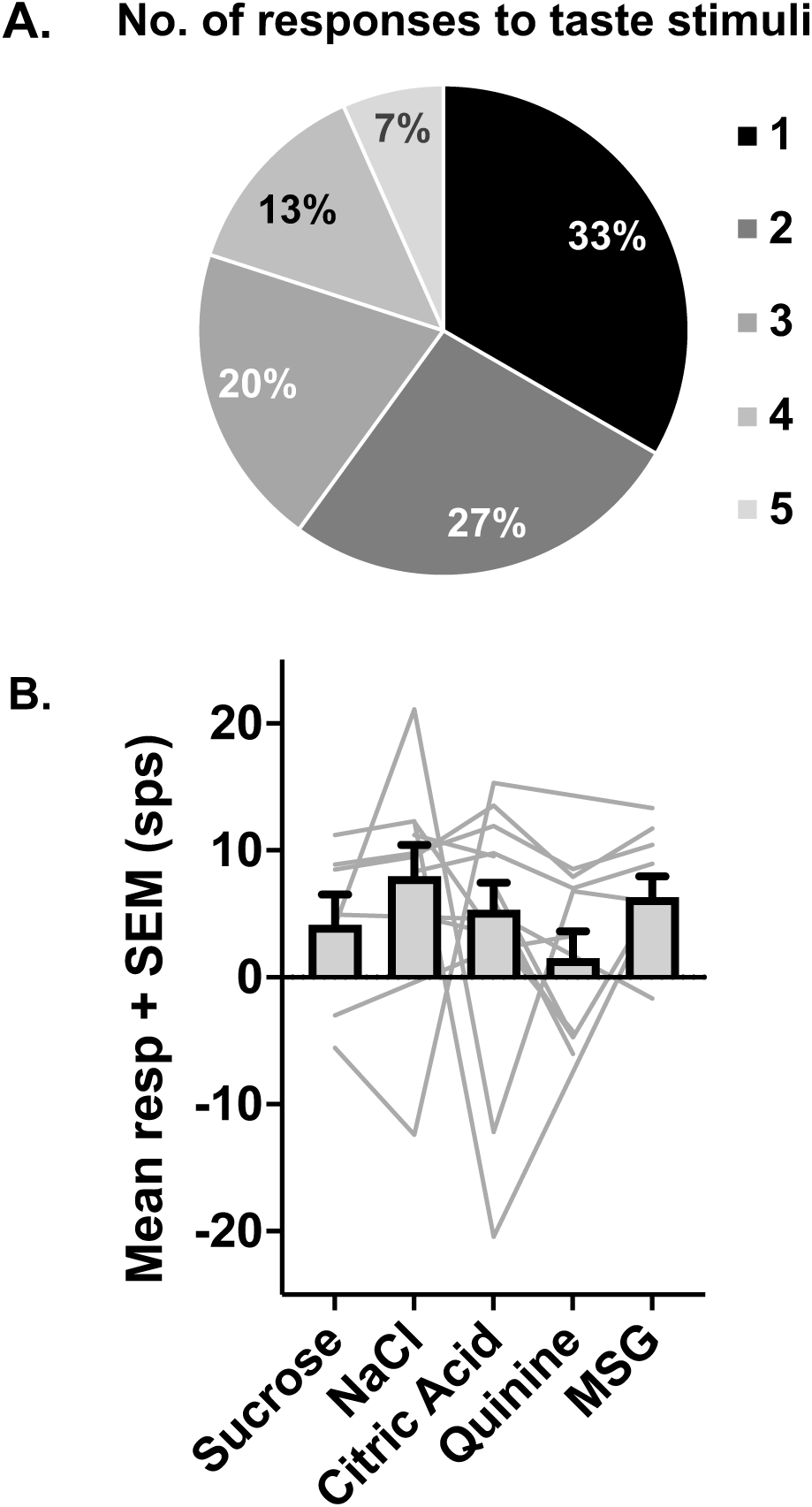
A. Proportion of taste-responsive cells that responded to 1-5 tastants tested in the Lick paradigm. Most cells (60%) responded to one or two tastants. B. Mean response rates (+SEM) evoked by each of the five tastants tested in the Lick paradigm. Light gray lines show the response profiles of individual cells. *N*= 28 cells.

**Figure 9.**
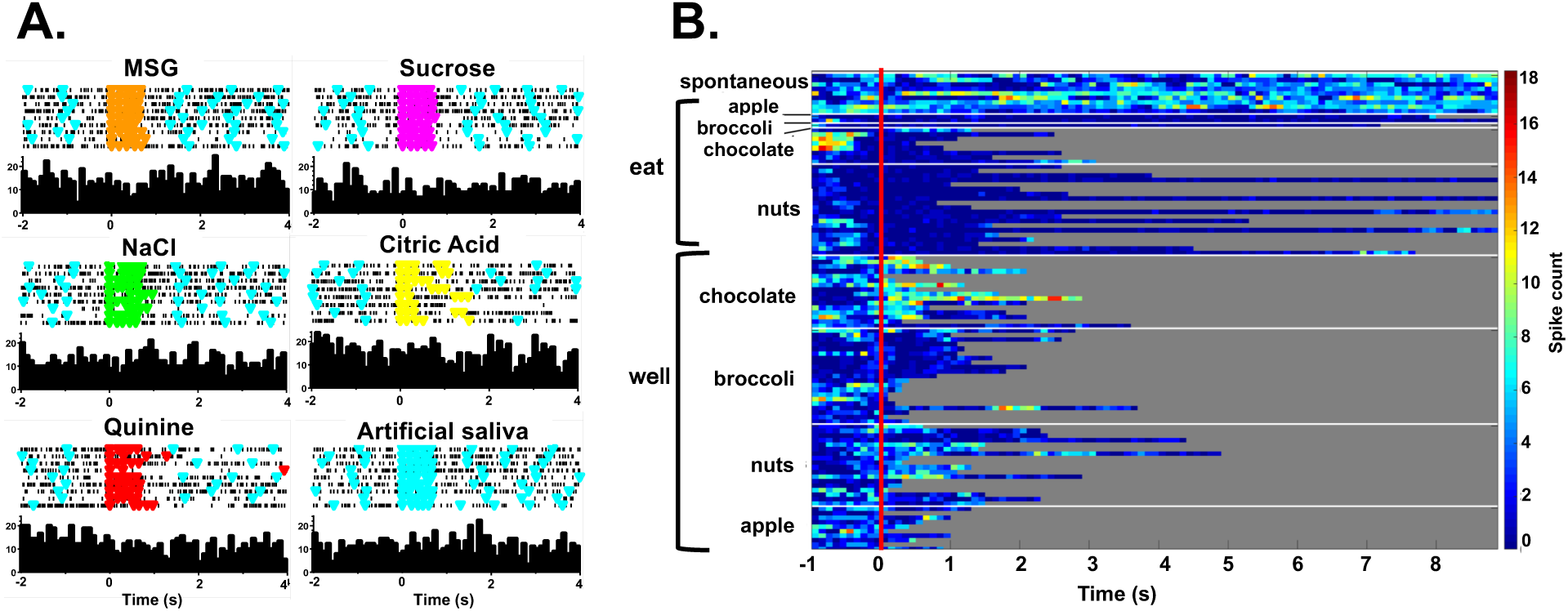
A. Peristimulus-time histograms (PSTHs) responses to tastants in one cell in the Lick paradigm. Top of each panel shows a raster plot where each line represents one trial. Bottom of each panel shows the resulting histogram of unit activity. This cell did not respond significantly to any tastant tested. B. Heat map of responses to well exploration and eating in the cell shown on the left. There was a robust response to well exploration for chocolate and a weaker response to nuts; however, the cell’s activity was below spontaneous firing rate during eating.

Although there was little response to the traditional taste stimuli, this cell during well entry for chocolate and nuts – but not during consumption of any food. In addition, there were many cells that were neither taste responsive nor responsive to food, but rather showed pronounced lick- related activity. Fig. 10 shows an example of this type of cell and its response to exploration of the food well and to eating. Surprisingly, this cell did not respond to eating, but showed a non- specific response to well exploration.

**Figure 10.**
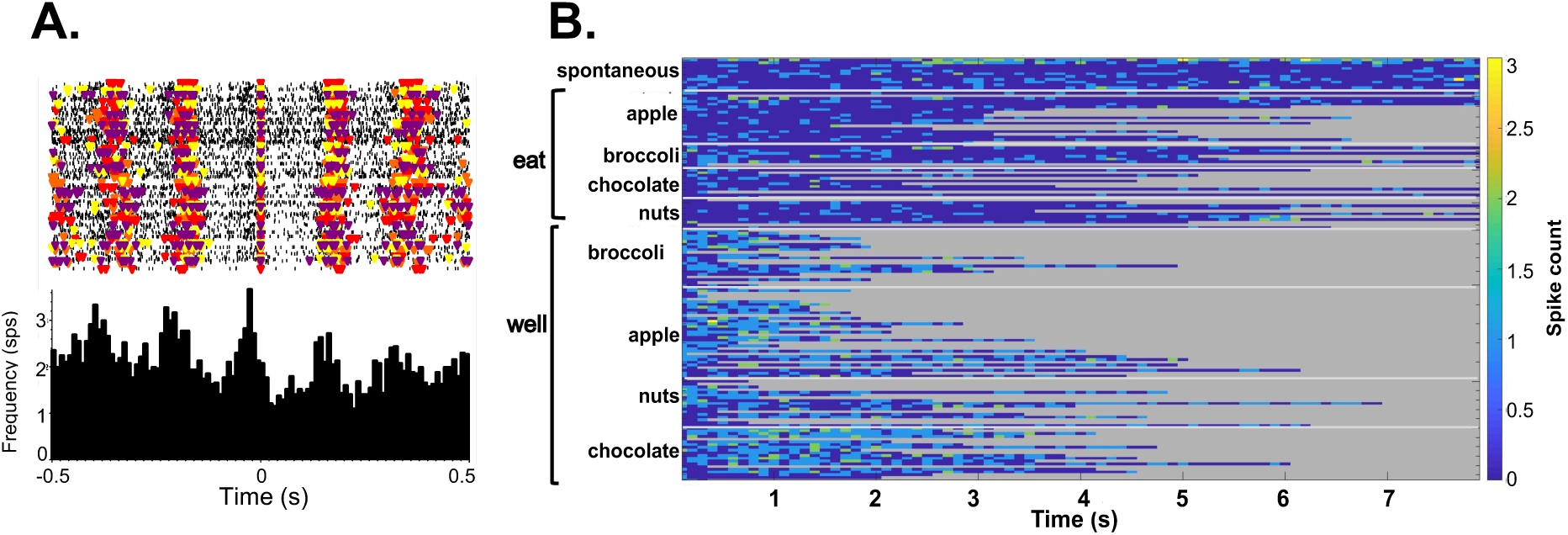
A. Raster (top) and PSTH (bottom) of activity in a lick-related cell. Each colored triangle on the raster plot denotes the occurrence of a lick; different colors show different tastants. On the PSTH, time zero denotes the occurrence of a lick. B. Heat map of the responses to well exploration and eating in the same cell shown in A. A weak, non-specific response to well exploration was evident in this cell; but the cell’s activity returned to spontaneous firing rates during eating.

Contrary to what one might expect, the response properties of each cell to the traditional taste stimuli tested in the Lick paradigm were not a good predictor of the responsiveness to the various foods. For example, we proposed that responses to sucrose would be associated with responses to milk chocolate, responses to NaCl would be associated with responses to salted peanuts, etc. There were too few responses to either taste stimuli or solid food in the sample of cells with responses to both tastants and solid food to calculate meaningful correlations.

However, even when cells were classified according to the stimulus that evoked their best, i.e., most vigorous, response, there was no sign of these predicted relationships (see Table 4). In addition, the food to which a cell responded best when exploring the food well was not a good predictor of the food to which a cell responded best when the food was consumed.

**Table 4.**
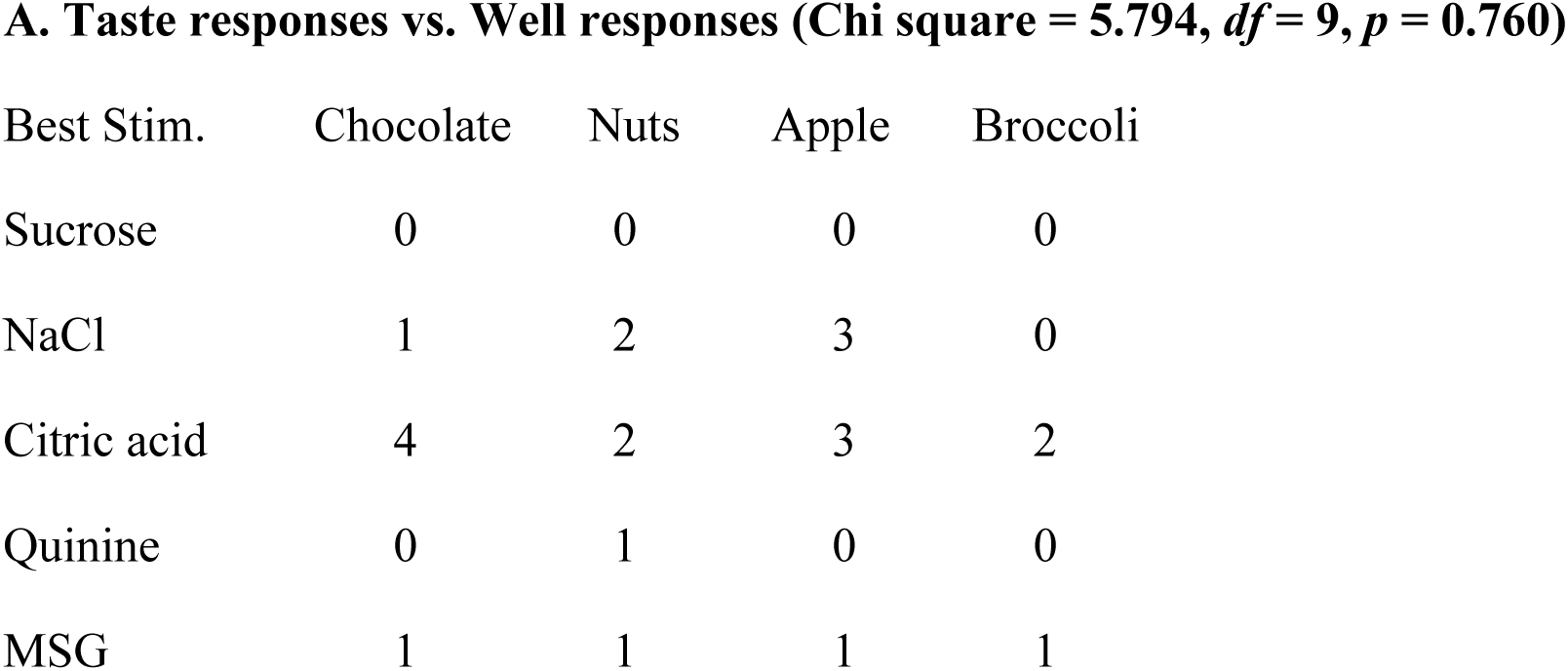
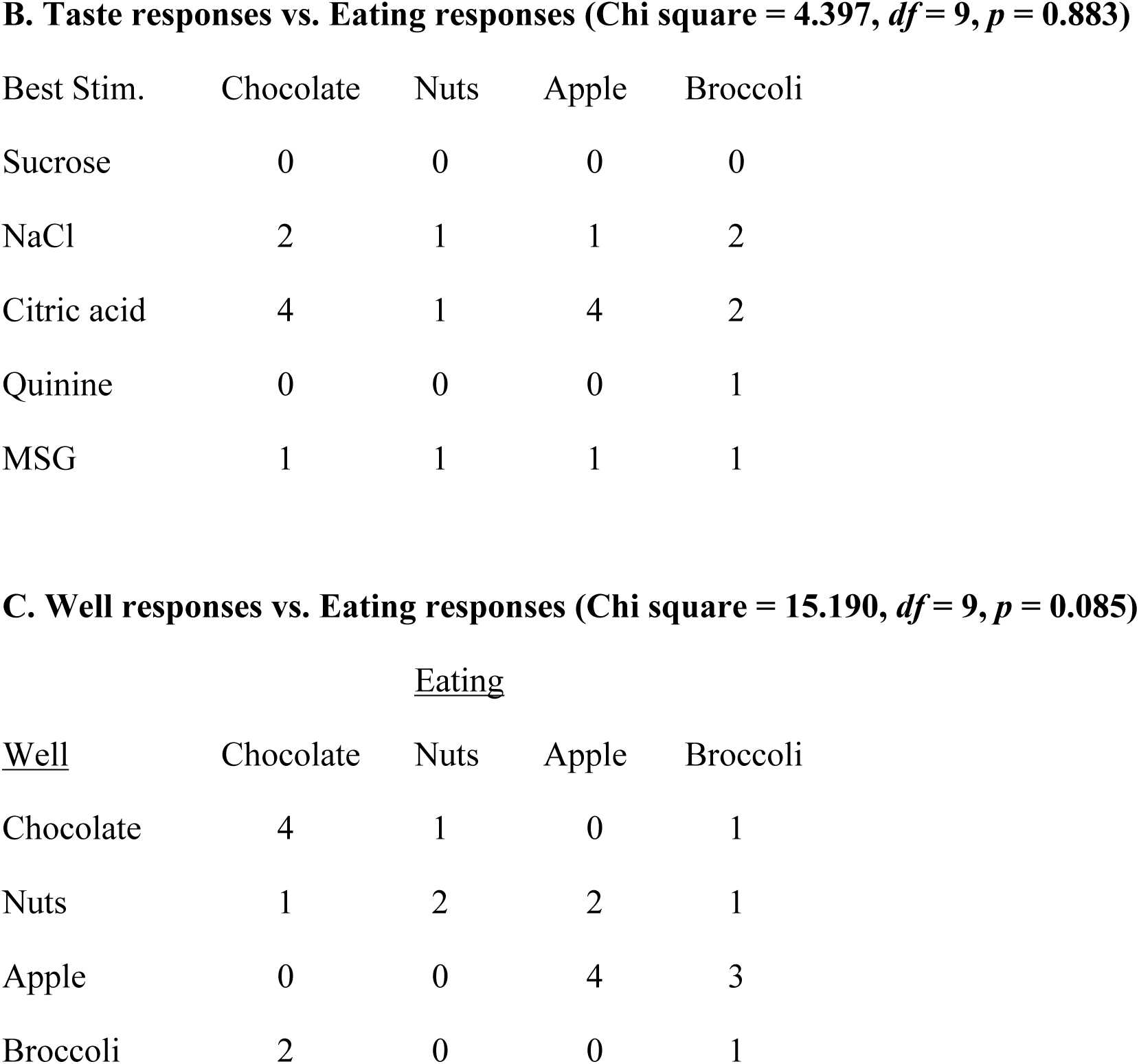
Relationship of taste responses (best stimulus) to responses to well and eating.

The data are publicly available at 10.6084/m9.figshare.25703352.

## Discussion

It has been frequently argued that the taste system is the final arbiter of ingestion; that is, its purpose is to identify sources of nutrition and avoid toxicity. However, the functionality of the taste system in the act of finding (appetitive) and ingesting (consummatory) solid food has rarely been studied (see Yamamoto et al., 1988, for an exception). Here, we recorded the responses of cells in the rNTS to licking traditional taste stimuli (Lick paradigm) and to performing appetitive and consummatory behaviors in a free feeding paradigm of solid foods (Eating). Results showed that the great majority of rNTS cells robustly and selectively increased their firing rate during exploration of the food-filled wells, often with responses beginning as a food well is approached but prior to any contact with the food. In contrast, most cells decreased their firing rate relative to responses to well entry during food consumption, at times showing firing rates significantly below spontaneous firing rates. Analyses of across neuron patterns of responsivity suggest that exploration and consumption evoke relatively independent response patterns. In addition, responses to tastants of various qualities in the Lick paradigm were poor predictors of responses to either well exploration or consumption of foods, even though these foods were chosen to have a single predominant taste quality. In fact, a lack of responsivity to liquid tastants did not preclude responsivity in the Eating paradigm, either for responsiveness to well exploration or food consumption. Collectively, these data suggest that a) both taste-responsive and non-taste-responsive cells in the rNTS preferentially transmit information about solid food in the appetitive phase of eating behavior, and b) by incorporating information from multiple sensory modalities, rNTS cells may convey more information relevant to evaluating food.

Nearly all cells in the rNTS responded more vigorously to the appetitive vs. the consummatory phases of eating, suggesting that there is a shift in how the population of cells carries the message for each function. Consistent with this idea are the results of MDS analyses of the across unit patterns of responses showing a clear separation of response patterns associated with well exploration vs. eating for all foods (see Fig. 6). Separate populations of cells for appetitive versus consummatory behaviors have been reported in the lateral hypothalamus (Jennings et al., 2015; Yamamoto et al., 1989) and arcuate nucleus (Mandelblat-Cerf et al., 2015), both of which have projections to the rNTS (Lundy and Norgren, 2004). Results presented here, however, do not support a similar arrangement in rNTS. That is, although the across unit patterns of response are clearly different, most rNTS cells respond to both well exploration and eating, albeit with generally weaker responses to eating. The attenuation of firing rate during eating in most cells is likely to reflect, at least in part, an active inhibition since many cells reduce their firing rate significantly below spontaneous firing rates while the animal eats. Such an inhibitory influence might arise from the considerable population of GABAergic cells in the rNTS (Boxwell et al., 2013; Davis, 1993; Lasiter and Kachele, 1988) or, alternatively, from centrifugal input (Smith and Li, 2000).

The anticipatory responses, which were common in NTS cells, were likely the result of both learning and olfactory responsiveness. Since these animals were very familiar with the experimental setup, a learned association between the odor and the taste of the food in the well may have developed over time, thus prompting an anticipatory response to contact with the food. It is possible that this activity originates in the brainstem given that cells in the rNTS are known to respond to olfactory as well as taste stimuli (Escanilla et al, 2015). However, since there are no known direct pathways from the olfactory bulb or piriform cortex to the rNTS, another possibility is that a centrifugal directive, potentially from the gustatory cortex (GC), triggers responses that precede well exploration. The gustatory cortex responds to olfactory stimuli (Samuelsen and Fontanini, 2017; Maier et al., 2015) and has direct projections to the rNTS (Lundy and Norgren, 2004). Relatedly, Gardner and Fontanini (2014) showed cue-related anticipatory responses to taste stimuli in the GC of alert mice. Moreover, Kusumoto-Yoshida et al. (2015) showed that the GC was not only critical for taste cue-related anticipatory activity, but that it also drove consummatory behavior.

### Licking vs. Eating

In the present experiment, both the liquid taste stimuli and the solid foods that were tested were chosen to span the gustatory domain of taste qualities, the so-called “basic” taste qualities: sweet, salty, sour, bitter and umami. Liquid taste stimuli were prototypical exemplars of each of the five basic taste qualities. Solid foods were chosen such that their predominant taste would correspond to one of the basic taste qualities. So, milk chocolate was chosen to represent sweetness, salted peanuts to represent saltiness, Granny Smith apples to represent sourness and broccoli to represent bitterness. The straightforward notion that sensitivity to the prototypical taste stimuli in the Lick paradigm would be reflected in the responses to the corresponding foods in the Eating paradigm, was not supported. In fact, there were mostly discrepancies between the most effective tastant in the Lick paradigm and the most effective food in the Eating paradigm (see Table 4). There are at least two factors that may have contributed to this mismatch. First, since the solid foods were each a combination of more than one taste quality, a one-to-one correspondence between responses to a prototypical liquid tastant and responses to the corresponding solid food would not be expected. Second, the concentrations of the prototypical liquid tastants may not have matched sufficiently the intensity of the taste qualities offered by the solid foods, and this may also tend to weaken the correspondence. However – what is striking is that we see no correspondence whatsoever, not even a weak one.

These two factors may also have contributed to the observation that even some non-taste responsive cells in the Lick paradigm nevertheless responded vigorously in the Eating paradigm. Alternatively, as Glendinning (2022) has argued, activity from the taste system alone may not be as informative about nutrition or toxicity as one might assume. Glendinning (2022) argues that input from other senses is necessary for a complete percept of a food. So, in that sense, it is not surprising that the taste system responds in one way when the animal licks sapid liquids and another way when faced with the richness of multisensory experiences provided by a solid food.

Surprisingly, the data show that taste-responsive cells in the rNTS participate preferentially in the appetitive vs. consummatory phase of ingestion. It is certainly predictable that the taste system is involved in the evaluative phase of eating; there is a large literature showing that taste-responsive cells in the rNTS respond selectively to tastants in the mouth. The surprising result reported here is that typical taste-responsive rNTS cells respond more vigorously to approach to a food with which they have had minimal or no contact, than to ingestion of the food. Cameras placed inside the food well in a few instances revealed that animals sniff and sample small bits of food accompanied by mouth and tongue movements.

These behaviors occur prior to grabbing larger chunks of food that they consume outside of the well. These data suggest that the behaviors that occur while the head is inside the well provide multimodal input (olfactory, somatosensory, gustatory, visual) to stimulate cells in the rNTS. In this regard, it is noteworthy that taste-responsive cells in the rNTS receive olfactory (Escanilla et al., 2015) as well as somatosensory input (Halsell et al., 1993; Hayma et al., 1985; Ogawa et al., 1984). Moreover, most cells in the rNTS show movement related activity as evidenced by the abundance of lick-related cells (Roussin et al., 2012; Denman et al. 2019). Surprisingly, lick-related cells in the present study responded to exploration of the food well, but not to eating *per se* (see Fig. 8). The convergence of information from several sensory modalities onto cells in the rNTS may enrich the information that can identify a food and inform the choice to consume or reject it.

The relatively subdued response of rNTS cells to eating compared with the robust response to well exploration was unexpected. One potential explanation is that the taste-related stimulation produced by mastication was not sufficient to drive a response. It is also possible that the information about food in the mouth while the animal is eating is processed by the caudal NTS, a region that is known to receive input from the vagus nerve innervating the gut (Kalia and Mesulam, 1980; Norgren and Smith, 1988). Thus, once the decision to consume a given food has been made based on input from the rNTS, the information flow then moves caudal so that the gut-brain pathways take over. In any case, the data suggest that the across unit patterns of response evoked by sensory input during the appetitive phase shift when the animal consumes the food.

## Acknowledgements

Supported by NIDCD Grant RO1-DC006914 to PMD

## References

Arif, Siti & Mohamad Mohsin, Mohamad Farhan & Abu Bakar, Azuraliza & Hamdan, Abdul & Syed Abdullah, Sharifah. (2017). Change point analysis: A statistical approach to detect potential abrupt change. Jurnal Teknologi. 79. 10.11113/jt.v79.10388.

Boxwell AJ, Yanagawa Y, Travers SP, Travers JB (2013) The mu-opioid receptor agonist DAMGO presynaptically suppresses solitary tract-evoked input to neurons in the rostral solitary nucleus. J Neurophysiol 109:2815–2826.

Chen Y, Lin YC, Kuo TW, Knight ZA (2015). Sensory detection of food rapidly modulates arcuate feeding circuits. Cell 160:829–841.

Cheng W, Gordian D, Ludwig MQ, Pers TH, Seeley RJ (2022) Hindbrain circuits in the control of eating behaviour and energy balance. Nature Metabolism, 4(7):826–835.

Davis BJ (1993) GABA-like immunoreactivity in the gustatory zone of the nucleus of the solitary tract in the hamster: light and electron microscopic studies. Brain Res Bull 30(1-2):69–77.

Denman AJ, Sammons JD, Victor JD, Di Lorenzo PM (2019) Heterogeneity of neuronal responses in the nucleus of the solitary tract suggests sensorimotor integration in the neural code for taste. J Neurophysiol. 121(2):634–645.

Di Lorenzo PM. (2021) Neural coding of food is a multisensory, sensorimotor function. Nutrients. 13(2):398.

Escanilla OD, Victor JD, Di Lorenzo PM (2015) Odor-taste convergence in the nucleus of the solitary tract of the awake freely licking rat. J Neurosci 35:6284–6297.

Gardner MP, Fontanini A (2014) Encoding and tracking of outcome-specific expectancy in the gustatory cortex of alert rats. J Neurosci 34(39):13000–17.

Glendinning JI. (2022) What does the taste system tell us about the nutritional composition and toxicity of foods? Handb Exp Pharmacol. 275:321–351.

Grill HJ, Hayes MR (2012) Hindbrain neurons as an essential hub in the neuroanatomically distributed control of energy balance. Cell Metab 16(3):296–309.

Halsell CB, Travers JB, Travers SP (1993) Gustatory and tactile stimulation of the posterior tongue activate overlapping but distinctive regions within the nucleus of the solitary tract. Brain Res 632:161–173.

Hayama T, Ito S, Ogawa H (1985) Responses of solitary tract nucleus neurons to taste and mechanical stimulations of the oral cavity in decerebrate rats. Exp. Brain Res. 60:235–242.

Hirata S, Nakamura T, Ifuku H and Ogawa H (2005) Gustatory coding in the precentral extension of area 3 in Japanese macaque monkeys; comparison with area G. Exp Brain Res 165:435–446.

Kalia M, Mesulam MM (1980) Brain stem projections of sensory and motor components of the vagus complex in the cat: I. The cervical vagus and nodose ganglion. J Comp Neurol 193:435– 465.

King MS (2007). Anatomy of the rostral nucleus of the solitary tract. In: The Role of the Nucleus of the Solitary Tract in Gustatory Processing (Bradley RM, ed), pp17–39. Boca Raton: Taylor and Francis.

Kusumoto-Yoshida I, Liu H, Chen BT, Fontanini A, Bonci A. (2015) Central role for the insular cortex in mediating conditioned responses to anticipatory cues. Proc Natl Acad Sci U S A. 112(4):1190–5.

Lasiter PS, Kachele DL (1988) Organization of GABA and GABA-transaminase containing neurons in the gustatory zone of the nucleus of the solitary tract. Brain Res Bull 21:623–636.

Lundy RF Jr., Norgren R (2004) Gustatory system. Inn: The Rat Nervous System (3rd edition) (G. Paxinos G, Mai J eds), pp891–921. San Diego: Academic Press.

Maier JX, Blankenship ML, Li JX, Katz DB. (2015) A multisensory network for olfactory processing. Curr Biol 25(20):2642–50.

Mandelblat-Cerf Y, Ramesh RN, Burgess CR, Patella P, Yang Z, Lowell BB, Andermann ML (2015) Arcuate hypothalamic AgRP and putative POMC neurons show opposite changes in spiking across multiple timescales. Elife. 4:e07122.

Norgren R, Smith GP (1988) Central distribution of subdiaphragmatic vagal branches in the rat. J Comp Neurol 273:207–223.

Ogawa H, Imoto T, Hayama T (1984) Responsiveness of solitario-parabrachial relay neurons to taste and mechanical stimulation applied to the oral cavity in rats. Exper Brain Res 54(2):349–358.

Rainer G, Asaad WF, Miller EK (1998) Selective representation of relevant information by neurons in the primate prefrontal cortex. Nature 393:577–579.

Roussin AT, D’Agostino AE, Fooden AM, Victor JD, Di Lorenzo PM (2012). Taste coding in the nucleus of the solitary tract of the awake, freely licking rat. J Neurosci 32(31):10494–10506.

Sammons JD, Weiss MS, Escanilla OD, Fooden AF, Victor JD, Di Lorenzo PM. Spontaneous changes in taste sensitivity of single units recorded over consecutive days in theb of the awake rat. (2016) PLoS One 11(8):e0160143.

Samuelsen CL, Fontanini A (2017). Processing of intraoral olfactory and gustatory signals in the gustatory cortex of awake rats. J Neurosci 37:244–257.

Schmitzer-Torbert N, Jackson J, Henze D, Harris K, Redish AD (2005) Quantitative measures of cluster quality for use in extracellular recordings. Neurosci 131(1):1–11.

Scott TR, Plata-Salaman CR (1999). Taste in the monkey cortex. Physiol Behav 67:489–511.

Smith DV, Li CS (2000) GABA-mediated corticofugal inhibition of taste-responsive neurons in the nucleus of the solitary tract. Brain Res 858:408–415.

Stevenson RJ, Prescott J, and Boakes RA (1995). The acquisition of taste properties by odors. Learn Motiv 26:1–23.

Travers S, Breza J, Harley J, Zhu J, Travers J (2018) Neurons with diverse phenotypes project from the caudal to the rostral nucleus of the solitary tract. J Comp Neurol. 526(14):2319–2338.

Tsang AH, Nuzzaci D, Darwish T, Samudrala H, Blouet C (2020) Nutrient sensing in the nucleus of the solitary tract mediates non-aversive suppression of feeding via inhibition of AgRP neurons. Mol Metab 42:101070.

Vincis R, Fontanini A (2016). Associative learning changes cross-modal representations in the gustatory cortex. eLife 5:e16420.

Vincis R, Fontanini A. (2019) Central taste anatomy and physiology. Handb Clin Neurol. 164:187–204.

Wilson DM, Lemon CH (2013) Modulation of central gustatory coding by temperature. J Neurophysiol 110:1117–1129.

Yamamoto T, Matsuo R, Kiyomitsu Y, Kitamura R (1988). Sensory inputs from the oral region in the cerebral cortex in behaving rats: An analysis of unit responses in cortical somatosensory and taste areas during ingestive behavior. J Neurophysiol 60(4):1303–1321.

Yarmolinsky DA, Zuker CS, Ryba NJP (2009) Common sense about taste: from mammals to insects. Cell 139:234–244.

